# Explainable models using transcription factor binding and epigenome patterns at promoters reveal disease-associated genes and their regulators in the context of cell-types

**DOI:** 10.1101/2024.05.06.592622

**Authors:** Omkar Chandra, Durjay Pramanik, Srishti Gautam, Madhu Sharma, Niharika Dubey, Biswarup Mahato, Yuriy L Orlov, Vibhor Kumar

## Abstract

Understanding genome-wide epigenetic regulation of diseases is important in establishing pathogenic factors and could aid in disease diagnosis, prognosis, and therapeutics. In this study, we have utilized transcription factors (TFs) and co-factor profiles (n=823) as features in machine learning models to link them to various diseases. Further, along with TFs and co-factor profiles, histone modifications ChIP-seq (n = 621), cap analysis gene expression (CAGE) tags (n = 255), and DNase hypersensitivity profiles (n = 255) as features allowed for the modeling of association of coding and non-coding genes to diseases. Such predicted associations could be independently validated using genome-wide association data and survival analysis. However, the unique aspect of our approach is that it highlights the link between TF binding patterns and diseases in the context of cell types. Besides highlighting relevant TF-binding in known cell-types associated with diseases, it also provided their surprising link with TFs expressed in immune cells and other seemingly non-related cells. Further investigation revealed such links to be genuine and potentially useful for prognosis, further revealing the need to deconvolve a set of known genes associated with diseases.

## Introduction

The expression of the genes involved in a given biological process is modulated by a set of few transcription and co-factors out of hundreds of transcription factors (TF) that bind at the regulatory sites of the genes. Dysregulation of genes involved in a biological process can lead to severe disease conditions. Researchers have sought to understand the role of epigenomics and transcription factors in different disease conditions. In a review, Van Ouwerkerk *et al*. discuss the role of epigenome and transcription factors in atrial fibrillation (AF) [1]. They describe that many of the AF-associated genomic loci lie within the non-coding regions, which are the binding sites of cardiac-specific transcription factors, having the causal role of affecting the gene expression by altering the activity of transcription factors and the epigenetic state of the chromatin. Mazzone *et al*. summarize the role of DNA methylation, histone modification, and non-coding RNAs in human autoimmune disorders like lupus, arthritis, and type 1 diabetes [2]. Therefore, Identifying epigenetic and transcription factor binding activities that are causal or drivers of dysregulated gene expression and genomic instability is an important task. Modeling the transcriptional regulatory mechanism of a disease can help reveal potential therapeutic targets and biomarkers for diseases.

Computational approaches have been used to identify key transcription factors in different phenotypic conditions. NetAct is a computational modeling approach to identify a core gene regulatory network of key transcription factors using expression data and a TF-target database. It utilizes gene co-expression patterns, cis-motif prediction, and TF-binding motif data [3]. Li *et al*. developed an integrated modeling framework using the time series expression data to identify the co-regulation of transcription factors, long non-coding RNAs (lncRNAs), and miRNAs on cardiac developmental dynamics. However, these approaches do not necessarily capture the direct binding of the transcription factors to the target genes, which is obtained from the ChIP-seq assay. The expression of a TF transcript does not necessarily mean the binding of the corresponding TF protein at the target site. Also, there is a rarity of studies on finding the link between diseases and TF binding patterns at the promoter sites of genes.

The objective of this study here is two-fold. First, we have modeled using the random forest algorithm to capture the association of different transcription factors and co-factors by using them as features across various disease gene-sets (DisGenNet). Second, We identified novel associations between genes (including non-coding) and diseases. The rationale behind this approach is that regulating both coding and non-coding genes depends on the active binding of multiple transcription factors and co-factors at the promoter region of these genes. The binding of the transcription factors depends upon the local genome’s open-chromatin state, which is reflected in the histone modifications and open chromatin conformation (ChIP-seq, DNase-seq) along with the expression of those genes. This same rationale is also true for the non-coding genes that are usually involved in regulating the coding genes.

Along with transcription factors and co-factors, epigenomic marks regulate the expression of coding and non-coding genes involved in the same biological process. It is possible to model this phenomenon to make novel associations between genes (coding and non-coding) and biological processes [4]. The identification of disease-associated genes has remained an important and challenging task. The associated genes can be casual or consequential genes exploited for therapeutic or prognostic purposes. The selection of candidate genes to link to a particular disease is a challenging task, which is the primary step before experimental evaluation of them. Especially linking non-coding RNA (ncRNA) with diseases has been recognized as an intriguing task due to several modes of action of ncRNA in regulating the coding genes [5,6].

Different computational methods use various strategies to associate genes with diseases. Some studies utilize the expression of the genes to link them to diseases [7–9]. However, most often, changes in the expression of genes due to diseases could be the consequence of the disease condition [10]. Therefore, associating genes with diseases merely due to changes in expression might not be reliable in identifying the causal genes. Some methods utilize gene-ontology knowledge as features to link more genes to diseases, but the drawback in using gene-ontology knowledge is that it does not accommodate for the prediction of the non-coding genes because the annotation of non-coding genes in the gene-ontology is very few (∼1200), therefore, modeling using the ontology knowledge graph-based suffers from sensitivity for non-coding genes. The protein-protein interactions feature does not allow for the prediction of non-coding genes. Finally, some approaches combine gene expression, protein-protein interactions, and GO knowledge of genes in the context of disease for predicting novel gene-disease links [11,12].

Most of these methods are merely prediction methods and can rarely provide regulatory insights about regulators of genes involved in disease or its phenotype. In addition, most machine learning-based methods are meant only for coding genes as they have annotations like biological processes and molecular functions. Hence, an important problem arises regarding predicting the association between non-coding genes and diseases to get regulatory inference. To address such problems, we have developed a novel method of associating coding and non-coding genes to diseases by utilizing transcription factor binding and epigenomic patterns. Our approach stands apart from other methods for predicting disease-gene association as it highlights regulatory aspects of TF and disease-associated genes and predicts non-coding genes involved in different disorders.

## Methods

### Epigenome and TF-binding features’ score calculation for promoters

We considered each disease gene-sets as a class and the annotated member genes as positives, and the association of novel genes is treated as a classification problem with epigenome and transcription binding assay data (DNAse-seq, ChIP-seq, CAGE tags) as features. The feature data used are of human samples and tissues aligned to the hg19 genome version. The ChIP-seq profiles of histone modification, open-chromatin profiles (DNAse-seq), and transcription factor binding for multiple human cell-type were downloaded from the ChIP-atlas database in bigWig format [13]. The bigWig files were converted to bedGraph file format using the bigWigToBedgraph tool. The scaled read-count scores for each genomic bin (200 bps) in the bedgraph files of ChIP-seq and CAGE-tag profiles were normalized by mean read-count for uniformity. For each ChIP-seq profile, the normalized read-count scores of genomic bins within 1 Kbp of transcription start sites (TSS) were added to get a read-score for a gene. Similarly, CAGE-tag profiles for multiple cell types were downloaded from the FANTOM (RIKEN, Japan) in bam file format, converted into BedGraph file format, and normalized the read-counts in the bins (200 bps) by mean read-count. The combined scores on bins lying within 1 kbp of TSS of a gene were combined to get scores for the gene.

The non-coding gene TSS was obtained from gencode (V30) and RefSeq gene transcripts [14,15]. Multiple TSS were allowed for each gene if their TSS were at least 500 bps away from each other. A total of 89747 promoter (genes) regions were analyzed.

### Prediction method

For each gene set in the DisGenNet, annotated genes were taken as positives, and an equal number of random genes were taken as negatives and disease-gene association prediction is treated as an interpretable classification problem using epigenome and TF binding data as features. Five machine learning models were trained for each gene set: random forest, XGBoost, SVM (support vector machine), linear regression-based lasso, and L2-regularization-based logistic regression (Ridge regression). Five-fold cross-validation was carried out to check the model’s performance for all the gene-sets. Model performance evaluation was carried out using metrics like accuracy, balanced accuracy, F1-score, Mathew’s correlation coefficient (MCC), and error rate. Linear regression function of ‘cv.glmnet’ was used with parameter alpha = 1). Logistic regression with L2-regularization (ridge regression model) was used with alpha = 0 and family = “binomial” from the ‘glmnet’ R package. For the random forests model, the ‘randomForest’ package with the ‘randomForest’ function.’ SVM was implemented using the ‘e1071’ R package with the ‘svm’ function. XGBoost models were implemented using the ‘xgboost’ R package with the ‘xgb.train’ function.

Novel associations of the genes to a disease gene-set were predicted using the trained models. Reliable scores were estimated by calculating the maximum precision for each gene-sets.

## Method for survival plots

Survival analysis was performed using the clinical and genomic information for patients of different cancer types in TCGA (The Cancer Genome Atlas). Related diseases were found using string matching between the names of the predicted diseases and TCGA cancer names. The ‘GEPIA2’ server [16] was used to acquire the survival plot of a gene of interest for one cancer. For any cancer, all the predicted genes of the associated disease (cancer-type) were combined and searched using the API of the server. After acquiring the survival plots for the predicted genes for cancer type, p(HR) values were extracted. p(HR) signifies the p-value of the ‘hazard ratio’ between high and low expression of a given gene for a cancer type. The hazard ratio can be considered a proxy for the extent of survival between two gene expression conditions (high and low). Python programming was used to implement the procedures. The ‘Null model’ was created for each cancer using the non-associated genes and random sampling. Survival plots and the p(HR) values were also acquired for the null hypothesis genes. Further, statistical analysis was performed on the p(HR) values between predicted genes and ‘null model’ genes for a cancer type. Mann–Whitney U test was used for the statistical analysis to find the significance (p-value < 0.05) of the predictions for a cancer type.

## Results

Each gene-set listed in the DisGenet [17] was treated as a class, and machine learning models were fitted using epigenome and TF ChIP-seq pattern at promoters of genes to predict their belongingness to each disease gene-set. We utilized all the features (epigenome, TF, co-factors, open-chromatin, and CAGE tag profiles) for gene-disease association prediction. For regulatory insights, we utilized just transcription factor binding as a feature.

### Epigenome and TF binding patterns at promoters are predictive of gene-disease association

Figure 1A shows the bar plots for the five-fold cross-validation result for 5 different machine-learning models, which indicates that the modeling is possible on disease gene sets using the TF binding and epigenome patterns at the promoters. Random forest models performed the highest compared to all the other models (linear regression, logistic regression, SVM, and XGBoost). The average AUROC from five-fold cross-validation for random forest models is 80%.

**Figure 1:**
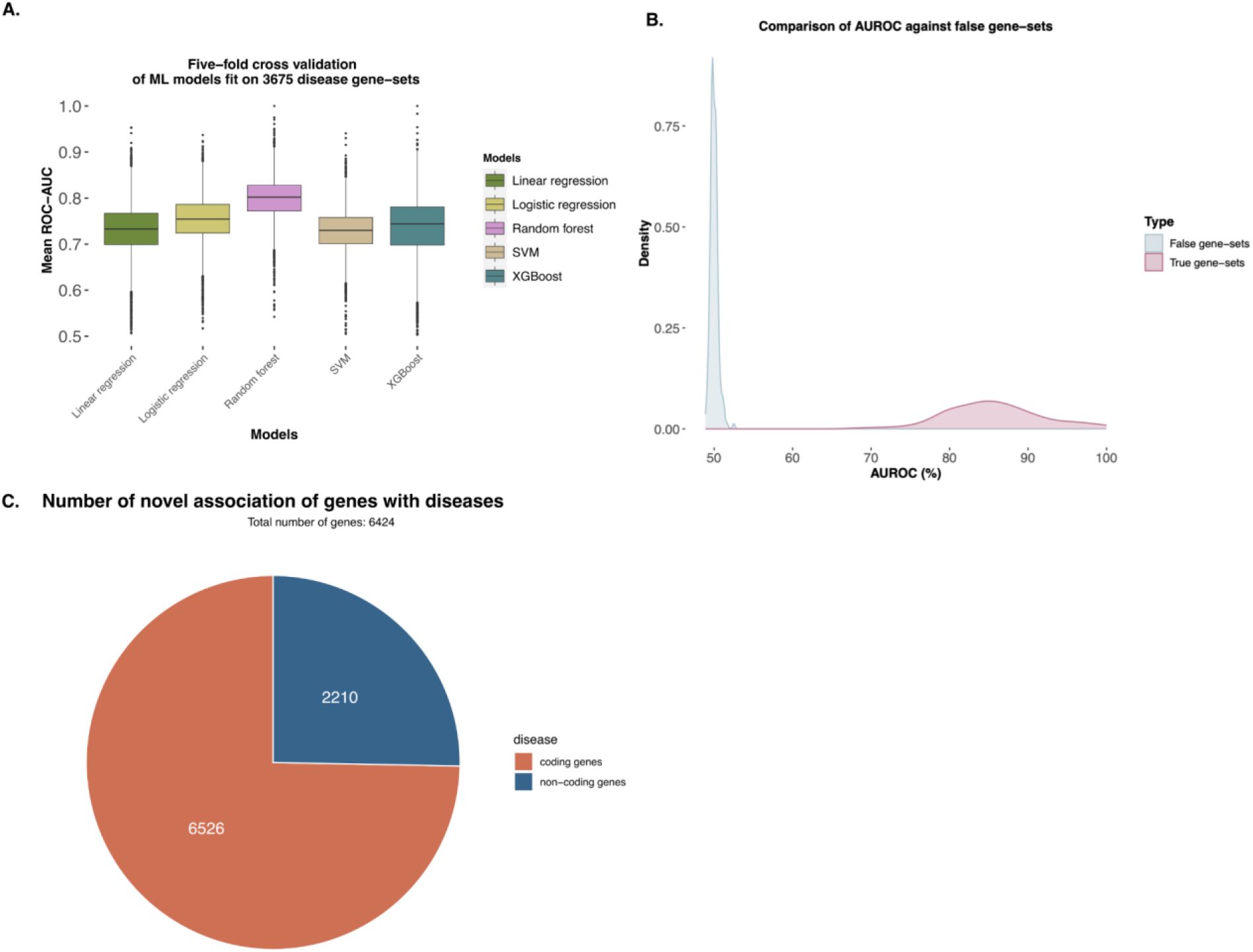
Results of prediction model used to find association between diseases and genes. A) Here 5 different machine-learning models were used to predict the membership of a gene in the gene-set for a disease. Five-fold cross-validation was done. The box plot shows the mean of ROC-AUC (Receiver operator characteristic-area under curve) from 5-five cross-validation for 3675 disease gene sets. B) The distribution of ROC-AUC achieved for original disease gene sets and false gene sets. C) Fraction of the number of coding and non-coding genes with predicted novel association with diseases using our approach.

### Specificity of the epigenome and TF binding as predictors of gene-disease association

To check if the modeling is non-random, the features of the genes were shuffled, the best performing ML algorithm, the random forest, was fit on the genes with random TF binding and epigenome signal at their promoters to check the performance, and the average AUROC was 55% for the null model (see Figure 1B), and for the same gene-sets with unshuffled features, the average AUROC was 85%. Such a result indicates that the TF binding and epigenome features are non-random and can be used to predict the belongingness of a gene to a disease gene set. Figure 1C shows the distribution of coding and non-coding genes associated with a disease.

### PubMed abstract mining-based validation

To validate the prediction of the association of the non-coding genes to the disease gene sets, an unbiased PubMed-based abstract mining was done to check the co-occurrence of the predicted non-coding gene term and the disease term in the abstracts published between 1990 and 2022. The co-occurrences of the predicted gene terms and disease terms are significantly more than the random non-coding gene and disease terms (see Figure 2A). This result indicates the reliability of predictions and gives confidence in predicting the association between non-coding genes and disease gene terms.

**Figure 2:**
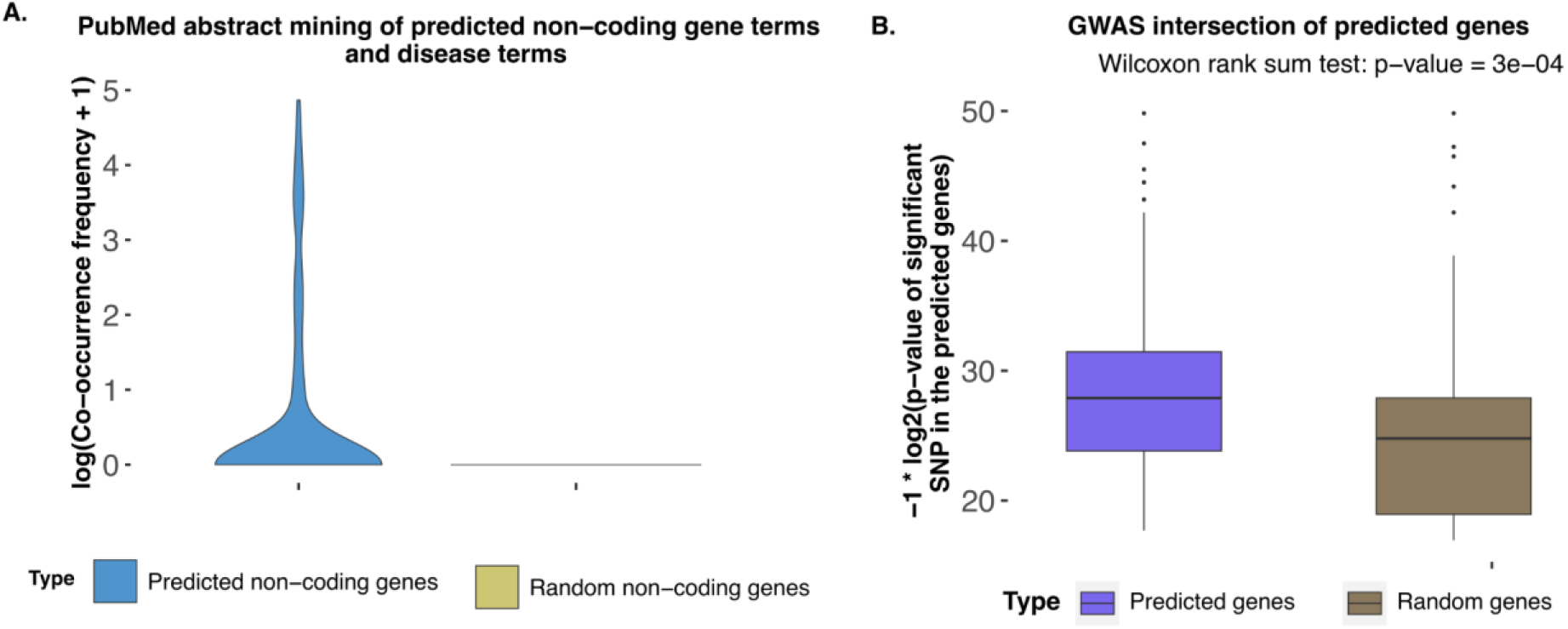
Validation of predicted associations between genes and diseases. A) validation based on PubMed abstract. Here, the Y-axis shows the frequency of co-occurrence of names of diseases and non-coding genes in the PubMed abstract. As a control, co-occurrence frequencies of random diseases and non-coding gene names in PubMed abstracts are also shown. B) The validation results using GWAS-based known mutations on genes predicted to be associated with diseases. Only GWAS mutation of same disease as predicted was counted. As a control, the number of GWAS mutations for target disease was counted on random genes.

### Validation using the result of GWAS

To further validate the predicted results, the presence of a mutation in the predicted genes for the same disease condition was checked using the GWAS database. The number of predicted genes that overlapped with the reported mutations was significantly more than the number of randomly associated genes and diseases (see Figure 2B)

### Survival analysis of predictive transcription factors and predicted genes for diseases

The features that rank at the top regarding feature importance while training and predicting the ML model are the transcription factors or DNA-binding proteins. These features are the components that were used for the ChIP-seq assay and the data of which was used for training and validation of the model on ground truth. A cutoff for feature importance value was fixed to find the set of TFs (or the genes associated with TFs) for each disease, referred to as top predictors.

Performing survival analysis with the top predictors for each cancer shows significant results in terms of hazard ratio. Figure 3 shows that for BLCA cancer, the significance between prediction and random selection of TFs is < 0.05. The subfigures suggest that some of the TFs from the pool of top BLCA predictors show promising results for cancer survival.

**Figure 3:**
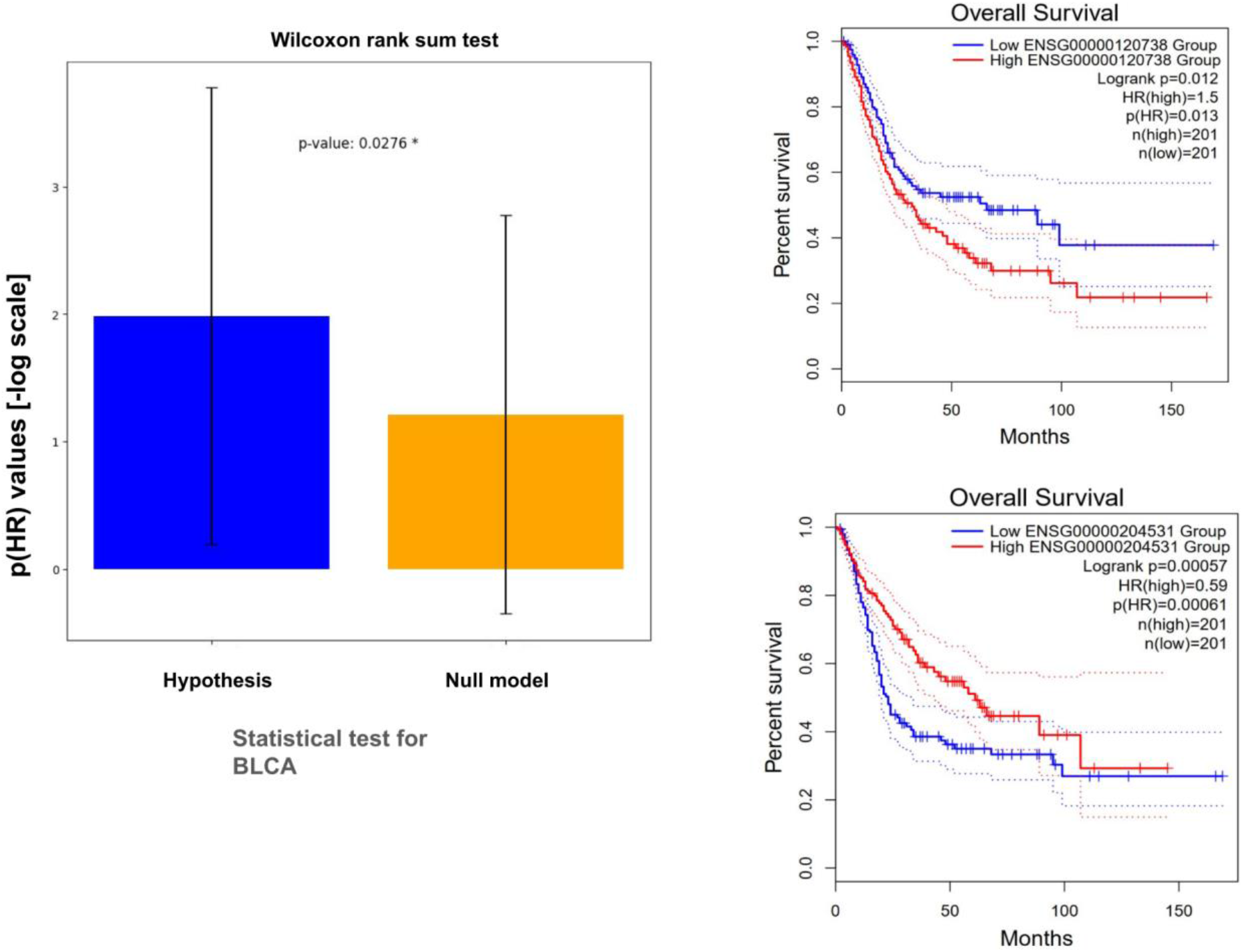
Association with survival for genes predicted to be linked with Bladder urothelial carcinoma (TCGA-BLCA). The left plot shows the box plot for -log(P-value) for association with survival, and the figures on the right show Kaplan-Meier plots for genes predicted to be associated with TCGA-BLCA.

In addition, the genes predicted for the cancers were also put through the survival analysis framework. However, the predicted genes (coding and non-coding) from those top predictors do not show promising results for cancer survival for high and low expression of the individual genes.

### Regulatory insights into the association between predictive TF and diseases

We could also model disease-gene association using TF and co-factors alone as features. See supplementary Figure 1A for model performance.

When such models were analyzed, the binding patterns of some TFs were top outlier predictors for diseases with good predictability. We focused on a few TF binding patterns in respective cell lines and found a few unknown associations between diseases and regulation by a few TFs (Supplementary Table 1). Results related to associations and related regulatory insights are shown below:

### Case study of ZNF366

We found that the promoter binding pattern of ZNF366 in dendritic cells appeared as a top outlier predictor for a few diseases, such as some types of lymphoma, Arthritis, Dermatitis, and even some types of Leukemia. ZNF366, also called DC-SCRIPT (dendritic cell-specific transcript), tends to have expressions specific to dendritic cells. Our results clearly indicated the role of dendritic cells in diseases where the binding pattern of ZNF366 (ChIP-seq in dendritic cells) appeared as the top predictor. For example, in Lymphomas (Adult, Childhood Diffuse Large B-cell, classical Hodgkin’s), there have been many studies that have shown the involvement of dendritic cells [18] [19]. Similarly, researchers have revealed the presence and role of defective dendritic cells in leukemia [20,21]. In addition to leukemia and lymphoma, ZNF366 appeared as the top predictor for genes associated with Dermatitis, pancreatitis, and encephalitis.

### Case study of LDB1

The binding pattern of a TF LDB1 (Lim Domain Binding protein1) in erythroblast cells was predictive for genes involved in anemia of two types, especially Hemolytic anemia and iron-refractory iron Deficiency anemia. Even though binding patterns of many TFs in many cell types were used as features, the appearance of the LDB1 binding pattern in erythroblast cells as the top predictor for anemia genes highlights our method’s specificity. LDB1 is one of the core master regulators involved in erythroid differentiation, and targeted deletion of Ldb1 in adult mice results in severe anemia [22]. Surprisingly, the LDB1 binding pattern in erythroblast appeared among the top outlier predictors for Apnea. Further detailed literature-based analysis revealed that the count and characteristics of erythrocytes could indeed be associated with Apnea [23][24]. We also found another disease, “Parkinson’s Disease, Familial, Type 1” (PDFT1), associated with LDB1 binding pattern in erythrocytes in our analysis. PDFT1, LDB1, and associated erythrocytosis appear to be causal, which is also partially supported by previous studies. Oligomeric α-synuclein in erythrocytes was elevated even in the initial motor stage of PD. For PDFT1, we also found promoter binding patterns of STAT5A in T Lymphocytes to be predictive of associated genes. Given the fact that neuroinflammation involved in the development of Parkinson’s disease is also mediated through T cells ([25] [26]), STAT5A in T cells seems to be an obvious regulator of genes associated with PDFT1. Thus, the top predictors for gene-sets disease not only hint about TFs but also cell types, which could contribute to the pathogenesis or symptoms of the diseases.

### Association of non-coding RNAs and disease

During the transcription event, the regulatory factors that govern the coding genes are also present across non-coding genes, especially for the genes involved in the same biological processes. Since the epigenome and transcription factor binding have been used in our disease-gene association models, it allowed the prediction of association between multiple non-coding genes to various diseases (see supplementary table 2).

Our model predicted that MIR137HG would be associated with the ‘Ductal Breast Carcinoma’ disease gene set. MIR137HG lies at chromosome 1, p21.3 position (see Figure 4A). MIR137HG was not in any gene set associated with breast cancer. Lee et al. investigated the role of MIR137HG and DEL1 genes using triple-negative breast cancer (TNBC) patient samples. Their luciferase reporter assay showed the direct binding of MIR137HG to the 3’-UTR of the DEL1 (developmental endothelial locus-1) gene. The DEL1 gene promotes tumor growth and apoptosis inhibition by altering the p53 pathway [27,28]. They have also shown that MIR137HG gene expression is downregulated, and DEL1 gene expression is upregulated compared to normal breast tissue samples. They concluded that TNBC cell proliferation, invasion, and migration were decreased after overexpressing the MIR137HG gene, which decreased the expression of the DEL1 gene [29].

**Figure 4:**
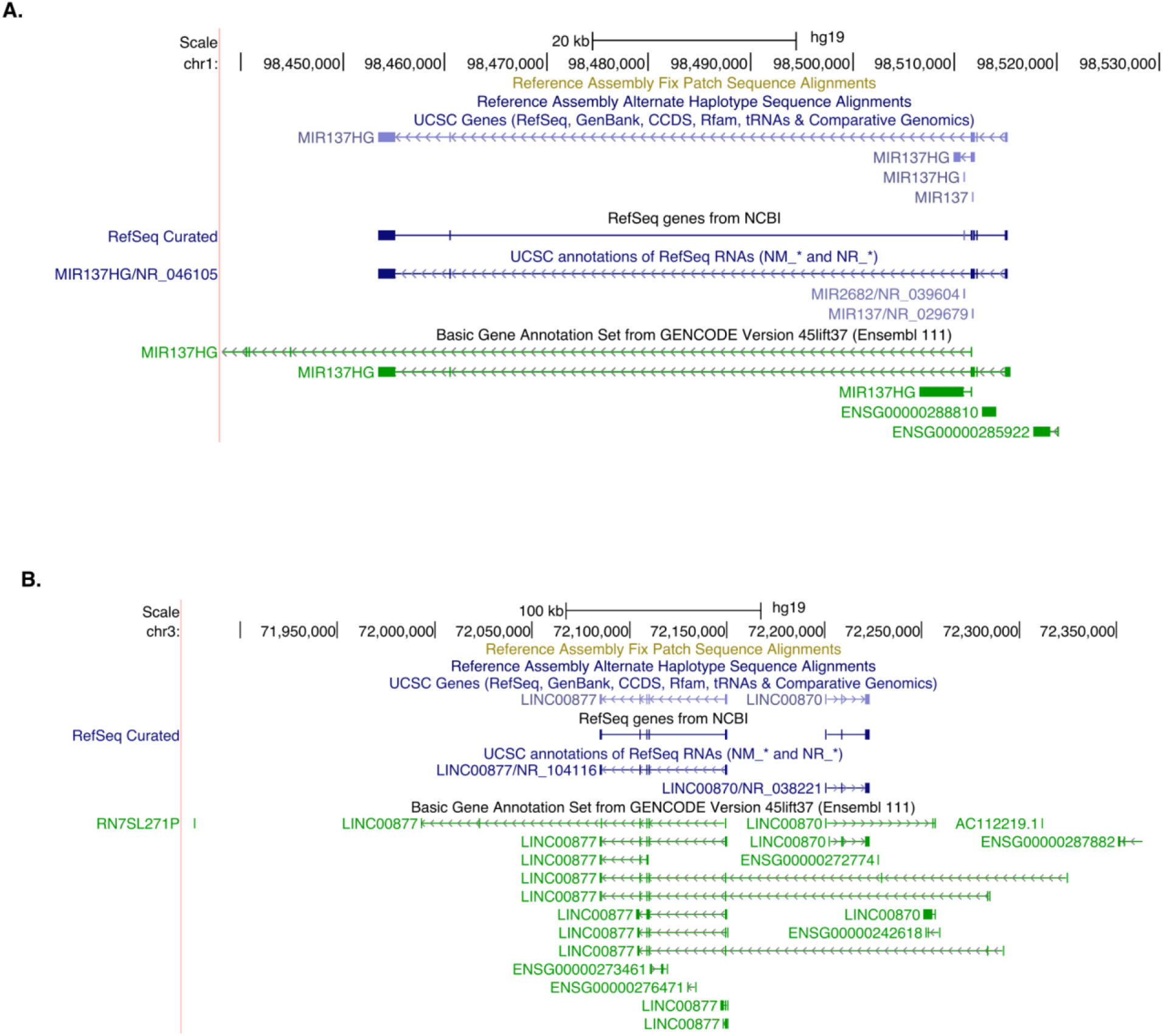
UCSC browser screenshot showing locations for two non-coding genes whose functions were predicted using our approach. A) location of gene MIR137HG B) LINC00877

Another long non-coding RNA, LINC00877, was found to be associated with the gene-set “Meningococcal Infections.” LINC00877 is present at chromosome 3, p13 band (see Figure 4B). Genome-wide association study results supported this association prediction; multiple variant and risk alleles of LINC00877 link to monocyte count (rs11708187-A, rs9809116-G, rs55890339-T). GWAS also links alleles of LINC00877 to platelet counts (rs9809116, rs11708187, rs12497693), which affects the development of the meningococcus prothrombotic trait [30]. Macrophages derived from monocytes have a prominent role in fighting infections, and platelet count indicates the severity of an infection [31].

## Discussion

Methods to find links between genes and diseases have been proposed by multiple groups. However, most of them focus on the accuracy of prediction and do not perform further analysis of important factors involved in regulating disease-associated genes. Rarely any other group has used epigenomic and transcription factor binding patterns to predict the relationship between diseases and genes, which allows finding links between non-coding RNAs and diseases. Moreover, no study based on regulatory inference rarely uses explainable prediction systems, such as those described here. Therefore, the uniqueness of our approach lies in the inference of top predictors as transcription factors, which leads to many genuine evidence-based insights. In addition to TFs, the cell-type for their ChIP-seq profiles also provided genuine insights into the involvement of different immune cell types in diseases. For example, a predicted association from our study links transcription factor ZNF366 ChIP-seq generated in a dendritic cell sample to psoriatic arthritis (see supplementary file 1). Marika *et al*. used the blood samples from generalized pustular psoriasis (GPP) patients to show that interleukin-36 production by immune cells acts directly on plasmacytoid dendritic cells, where it potentiates toll-like receptor (TLR)-9 activation and IFN-α production [32]. Increased IFN-α triggers feed-forward inflammation, causing increased psoriatic extracutaneous comorbidities. Studies have shown that individuals with such conditions have a higher risk of developing psoriatic arthritis [33]. Marika *et al*. have also shown that the ZNF366 gene is overexpressed in the blood of GPP patients. Another example from our analysis, ZNF366 ChIP-seq generated from a dendritic cell sample, was linked to ‘acute pancreatitis’ (see supplementary file 1). Bedrosian et al. have shown that dendritic cells are important for pancreas viability and may protect the pancreas after an episode of pancreatitis in mice [34]. They also showed that pancreatic dendritic cells produced markedly elevated levels of activating cytokines like IFN-γ, and it has been shown that transcription factor DC-SCRIPT encoded by ZNF366 regulates the cytokines IFN-γ and IL-12p40 levels in dendritic cells [34,35]. Thus, our approach can provide inferences to link different cell types to diseases and help researchers develop novel hypotheses.

One important result of our study is predicting the link between non-coding genes and diseases. (supplementary file2). A few examples are as follows: a) CDKN2B-AS1 and ‘Pancreatic Ductal Carcinoma,’ a study by Giaccherini et al. reported that rs1412832 polymorphism, that is situated in the CDKN2B-AS1/ANRIL, showed a genome-wide significant association with increased risk of developing pancreatic ductal adenocarcinoma [36]. b) ATXN8OS and ‘drug response’, Luo et al. conducted a study using glioma xenograft models to show that ATXN8OS inhibited temozolomide (TMZ)-resistance of glioma [37]. As such, much more evidence from the literature supports the predictions of our model (see supplementary file 2).

As utilizing an explainable model for highlighting the importance of TF and their binding profile in various cell-types was one of the main purposes of this study, we did not perform benchmarking to compare the accuracy of prediction of disease-gene association. Through our analysis, we highlighted many genes and TFs that could be associated with different types of cancer. For example, our downstream analysis linked TF RELB from the ChIP-seq GM12878 cell line to primary peritoneal carcinoma (see Supplementary File 1). Marcela A. et al., in their work, report that B cells localized within the peritoneal cavity produce IgM and protect against tumor growth [38], and it has been found that the RELB gene is essential for maintaining the B cell development and maturation [39]. Another example from our analysis is the linkage between TF CREB1 from ChIP-seq GM12878 cell line and ontology class, “Tumor Promotion” (see supplementary file1). Yang et al. have shown that B-cells promote tumor progression through angiogenesis in melanoma and lung carcinoma, and Frissora et al. have shown that CREB-1 binding at CRE (cyclic-AMP response elements) induces B-cell proliferation. Thus, our downstream helps link immune cell types to cancer [40,41]. It generated many putative hypotheses. We validated many putative hypotheses of association between top predictive TF and cancer using survival analysis with TCGA-based clinical data sets.

An important trend that can be noticed from associations based on predictive TFs and diseases is that it shows divergence and convergence of cell types involved in diseases. Often, genome-wide association studies (GWAS) based association between genes and disease does not reveal the relevance of any cell-type. Due to the inherent property of our method to use the context of cell-types with promoter binding profile of TF, it has vast potential to guide validations in relevant cell-types. Highlighting cell-types can also impact diagnostic, prognostic, and therapeutic decisions. As highlighted by our study, the genes associated with Apnea and bound by the top predictive TF (see supplementary file1) for Apnea have expression and functional activity in erythroblast. It is well known that the human body produces more erythrocytes to cope with low oxygen levels due to apnea. Thus, erythrocyte properties and count can help the prognosis of sleep Apnea [24]. However, the activity of those genes themselves could be causal or an effect of apnea. On a similar pattern, our results highlight the involvement of different immune cell types in many disease conditions. Thus, the current annotation of the ontology consists of genes captured from the DE analysis using bulk RNA expression data. The bulk samples could consist of local tissue-specific cells and immune cells.

Many examples hint that the current set of genes associated with diseases could consist of genes active in immune cells but get dysregulated due to disease (effect). However, the gene-set for a disease could also consist of causal genes active in the immune cell. Our analysis has the potential to highlight both aspects to some extent, which hints towards a broader fundamental problem with the current gene-set associated with diseases. There is no denying that genes affected by a disorder can be used for diagnosis or prognosis; however, there is a need to fine-tune the annotation and deconvolve the causality and effect of the members of gene sets currently known to be associated with diseases.

## References

1. van Ouwerkerk AF, Hall AW, Kadow ZA, Lazarevic S, Reyat JS, Tucker NR, et al. Epigenetic and Transcriptional Networks Underlying Atrial Fibrillation. Circ Res. 2020 [cited 8 Feb 2024]. doi:10.1161/CIRCRESAHA.120.316574

2. Mazzone R, Zwergel C, Artico M, Taurone S, Ralli M, Greco A, et al. The emerging role of epigenetics in human autoimmune disorders. Clin Epigenetics. 2019;11: 1–15.

3. Su K, Katebi A, Kohar V, Clauss B, Gordin D, Qin ZS, et al. NetAct: a computational platform to construct core transcription factor regulatory networks using gene activity. Genome Biol. 2022;23: 1–21.

4. Patterns of transcription factor binding and epigenome at promoters allow interpretable predictability of multiple functions of non-coding and coding genes. Comput Struct Biotechnol J. 2023;21: 3590–3603.

5. Kaikkonen MU, Lam MTY, Glass CK. Editor’s Choice: Non-coding RNAs as regulators of gene expression and epigenetics. Cardiovasc Res. 2011;90: 430.

6. Patil VS, Zhou R, Rana TM. Gene regulation by noncoding RNAs. Crit Rev Biochem Mol Biol. 2014;49: 16.

7. Qumsiyeh E, Showe L, Yousef M. GediNET for discovering gene associations across diseases using knowledge based machine learning approach. Sci Rep. 2022;12: 1–17.

8. Shah SD, Braun R. GeneSurrounder: network-based identification of disease genes in expression data. BMC Bioinformatics. 2019;20: 1–12.

9. Nam Y, Jhee JH, Cho J, Lee J-H, Shin H. Disease gene identification based on generic and disease-specific genome networks. Bioinformatics. 2018;35: 1923–1930.

10. Porcu E, Sadler MC, Lepik K, Auwerx C, Wood AR, Weihs A, et al. Differentially expressed genes reflect disease-induced rather than disease-causing changes in the transcriptome. Nat Commun. 2021;12: 1–9.

11. Asif M, Martiniano HFMC, Vicente AM, Couto FM. Identifying disease genes using machine learning and gene functional similarities, assessed through Gene Ontology. PLoS One. 2018;13. doi:10.1371/journal.pone.0208626

12. Chen B, Wang J, Li M, Wu F-X. Identifying disease genes by integrating multiple data sources. BMC Med Genomics. 2014;7: 1–12.

13. Oki S, Ohta T, Shioi G, Hatanaka H, Ogasawara O, Okuda Y, et al. ChIP-Atlas: a data-mining suite powered by full integration of public ChIP-seq data. EMBO Rep. 2018;19. doi:10.15252/embr.201846255

14. Frankish A, Diekhans M, Jungreis I, Lagarde J, Loveland JE, Mudge JM, et al. GENCODE 2021. Nucleic Acids Res. 2021;49: D916–D923.

15. O’Leary NA, Wright MW, Brister JR, Ciufo S, Haddad D, McVeigh R, et al. Reference sequence (RefSeq) database at NCBI: current status, taxonomic expansion, and functional annotation. Nucleic Acids Res. 2016;44: D733–45.

16. Tang Z, Kang B, Li C, Chen T, Zhang Z. GEPIA2: an enhanced web server for large-scale expression profiling and interactive analysis. Nucleic Acids Res. 2019;47: W556–W560.

17. Piñero J, Bravo À, Queralt-Rosinach N, Gutiérrez-Sacristán A, Deu-Pons J, Centeno E, et al. DisGeNET: a comprehensive platform integrating information on human disease-associated genes and variants. Nucleic Acids Res. 2016;45: D833–D839.

18. Bhardwaj N, Brody JD. Dendritic cells and lymphoma cells: come together right now. Blood. 2015;125: 5–7.

19. Kim SH, Choe J-Y, Jeon Y, Huh J, Jung HR, Choi Y-D, et al. Frequent expression of follicular dendritic cell markers in Hodgkin lymphoma and anaplastic large cell lymphoma. J Clin Pathol. 2013;66: 589–596.

20. Circulating blood dendritic cells from myeloid leukemia patients display quantitative and cytogenetic abnormalities as well as functional impairment. Blood. 2001;98: 3750–3756.

21. Mohty M, Jarrossay D, Lafage-Pochitaloff M, Zandotti C, Brière F, de Lamballeri XN, et al. Circulating blood dendritic cells from myeloid leukemia patients display quantitative and cytogenetic abnormalities as well as functional impairment. Blood. 2001;98. doi:10.1182/blood.v98.13.3750

22. Hattangadi SM, Wong P, Zhang L, Flygare J, Lodish HF. From stem cell to red cell: regulation of erythropoiesis at multiple levels by multiple proteins, RNAs, and chromatin modifications. Blood. 2011;118: 6258–6268.

23. Increased erythrocyte adhesiveness and aggregation in obstructive sleep apnea syndrome. Thromb Res. 2008;121: 631–636.

24. Narváez PA, Mohrenberger CJ, Baena EM, Rivera CG, Villalona RM, Meneses PL, et al. Erythrocytosis in patients with obstructive sleep apnea. Eur Respir J. 2014;44. Available: https://erj.ersjournals.com/content/44/Suppl_58/P2210.abstract

25. The role of T cells in the pathogenesis of Parkinson’s disease. Prog Neurobiol. 2018;169: 1–23.

26. Contaldi E, Magistrelli L, Comi C. T Lymphocytes in Parkinson’s Disease. J Parkinsons Dis. 2022;12: S65.

27. Kim H, Lee SH, Lee MN, Oh GT, Choi KC, Choi EY. p53 regulates the transcription of the anti-inflammatory molecule developmental endothelial locus-1 (Del-1). Oncotarget. 2013;4. doi:10.18632/oncotarget.1318

28. Lee SH, Kim DY, Jing F, Kim H, Yun CO, Han DJ, et al. Del-1 overexpression potentiates lung cancer cell proliferation and invasion. Biochem Biophys Res Commun. 2015;468. doi:10.1016/j.bbrc.2015.10.159

29. Lee SJ, Jeong J-H, Kang SH, Kang J, Kim EA, Lee J, et al. MicroRNA-137 Inhibits Cancer Progression by Targeting Del-1 in Triple-Negative Breast Cancer Cells. Int J Mol Sci. 2019;20. doi:10.3390/ijms20246162

30. Lécuyer H, Borgel D, Nassif X, Coureuil M. Pathogenesis of meningococcal purpura fulminans. Pathog Dis. 2017;75: ftx027.

31. Peiser L, De Winther MP, Makepeace K, Hollinshead M, Coull P, Plested J, et al. The class A macrophage scavenger receptor is a major pattern recognition receptor for Neisseria meningitidis which is independent of lipopolysaccharide and not required for secretory responses. Infect Immun. 2002;70. doi:10.1128/IAI.70.10.5346-5354.2002

32. IL-36 Promotes Systemic IFN-I Responses in Severe Forms of Psoriasis. J Invest Dermatol. 2020;140: 816–826.e3.

33. David Burden A, Kirby B. Psoriasis and Related Disorders. Rook’s Textbook of Dermatology, Ninth Edition. John Wiley & Sons, Ltd; 2016. pp. 1–64.

34. Bedrosian AS, Nguyen AH, Hackman M, Connolly MK, Malhotra A, Ibrahim J, et al. Dendritic Cells Promote Pancreatic Viability in Mice with Acute Pancreatitis. Gastroenterology. 2011;141: 1915.

35. Zhang S, Coughlan HD, Ashayeripanah M, Seizova S, Kueh AJ, Brown DV, et al. Type 1 conventional dendritic cell fate and function are controlled by DC-SCRIPT. Science Immunology. 2021 [cited 21 Feb 2024]. doi:10.1126/sciimmunol.abf4432

36. Giaccherini M, Farinella R, Gentiluomo M, Mohelnikova-Duchonova B, Kauffmann EF, Palmeri M, et al. Association between a polymorphic variant in the CDKN2B-AS1/ANRIL gene and pancreatic cancer risk. Int J Cancer. 2023;153: 373–379.

37. Luo J, Bai R, Liu Y, Bi H, Shi X, Qu C. Long non-coding RNA ATXN8OS promotes ferroptosis and inhibits the temozolomide-resistance of gliomas through the ADAR/GLS2 pathway. Brain Res Bull. 2022;186. doi:10.1016/j.brainresbull.2022.04.005

38. Haro MA, Dyevoich AM, Phipps JP, Haas KM. Activation of B-1 Cells Promotes Tumor Cell Killing in the Peritoneal Cavity. Cancer Res. 2019;79: 159–170.

39. Laidlaw BJ, Cyster JG. Transcriptional regulation of memory B cell differentiation. Nat Rev Immunol. 2020;21: 209–220.

40. Yang C, Lee H, Pal S, Jove V, Deng J, Zhang W, et al. B Cells Promote Tumor Progression via STAT3 Regulated-Angiogenesis. PLoS One. 2013;8: e64159.

41. Frissora F, Chen H-C, Durbin J, Bondada S, Muthusamy N. IFN-γ-mediated inhibition of antigen receptor-induced B cell proliferation and CREB-1 binding activity requires STAT-1 transcription factor. Eur J Immunol. 2003;33: 907–912.

